# Posture controls mechanical tuning in the black widow spider mechanosensory system

**DOI:** 10.1101/484238

**Authors:** Natasha Mhatre, Senthurran Sivalinghem, Andrew C. Mason

## Abstract

Spiders rely on mechanical vibration sensing for sexual signalling, prey capture and predator evasion. The sensory organs underlying vibration detection, called slit sensilla, resemble cracks in the spider’s exoskeleton, and are distributed all over the spider body. Those crucial to sensing web- and other substrate-borne vibrations are called lyriform organs and are densely distributed around leg joints. It has been shown that forces that cause bending at leg joints also activate these lyriform organs. Little is known of how the biomechanics of the body of a freely-suspended spider in its natural posture interact with vibrations introduced into the body and how this affects vibration perception. Female black widow spiders, in particular, have a striking body-form; their long thin legs support a large pendulous abdomen. Here, we show that in their natural posture, the large abdominal mass of black widow females, interacts with the spring-like behaviour of their leg joints and determines the mechanical behaviour of different leg joints. Furthermore, we find that adopting different body postures enables females to alter both the level and tuning of the mechanical input to lyriform organs. Therefore, we suggest that posture may be used to flexibly and reversibly focus attention to different classes or components of web vibration. Postural effects thus emphasize the dynamic loop of interactions between behaviour and perception, i.e. between ‘brain’ and body.

## Introduction

Vibration sensing is crucial to spiders, to perceive, and localize predators, prey, and mates(1). The sensory organs that enable vibration sensing are called slit sensilla and they function by sensing minute strains in the exoskeleton(2). The slit sensilla that are most important in the context of detecting substrate-borne vibrations are grouped into Lyriform Organs - compound organs composed of several slits closely arranged in parallel and densely distributed near leg joints(1). Much of what is known about lyriform organs is known through studying neurophysiology. Neurophysiological studies in spiders have carefully concentrated on individual legs, joints, and/or receptors and made highly precise measurements by delivering vibrational stimuli to these individual joints (3–6).Extremely detailed work over several decades has shown that bending a leg joint is sufficient to activate lyriform organs and to produce neurophysiological responses(1, 7–9). The exact physical mechanism that converts leg bending into the exoskeletal strains that excite neuronal responses is not completely understood. In general, it is believed that strains are transmitted to the exoskeleton during joint bending through some combination of local hydrostatic pressure changes in the spider’s legs, and through direct mechanical ligament attachments(4, 7, 10, 11). Indeed, lyriform organs are exquisitely sensitive(3–5) and can sense very small vibrations propagating through the substrate or the web(6, 12). All lyriform organs studied to date show broad frequency tuning responding equally well to most frequencies observed in substrate-borne vibrations(3, 4). It is only further upstream, in the central nervous system, that some degree of frequency discrimination is observed(9).

Before the nervous system becomes involved, however, the mechanical input to the lyriform organs is first shaped by the mechanics of the spider’s body and its environment. Older measurements, which were often made using extremely innovative measurement systems made by the authors themselves, have shown that web tension and the coupling of the leg to web have strong effects on the vibrations delivered to the leg(6). Even older work had indicated that the mechanics of the spider’s body could modulate the vibrations delivered to the joints (13, 14). It is increasingly recognized that perception is a cognitive process that occurs through the interaction of the nervous system, the body, and the environment in which the body is situated (15, 16). Even animals with relatively simple nervous systems, like spiders, can act as adaptive, problem-solving agents that use their bodies and environment to modulate perception, thus demonstrating embodied or extended cognition(17, 18). As a first step towards a more complete picture of spider vibration perception, we decided to exploit the sensitivity of modern measurement techniques and the power of current modelling tools to test how the mechanics of the spider’s body, arising from its natural posture on the web would affect its perception (3, 4).

In web-dwelling spiders, vibration perception becomes particularly rich and interesting given that both the environment and the body can be manipulated by the spider (17, 19–21). Spiders effectively develop and alter their environment to suit their perceptual needs, *via* their web (21–23). A great deal of research now has considered the effect and contribution of the web as an extension of spider perception (22, 24–27). The role of the other player in the system: the reconfigurable, dynamic, and flexible spider body has, with a few exceptions (4, 6), remained largely unstudied. Here we explore the role of the body of the female black-widow spider (*Latrodectus hesperus*) in vibration perception. Specifically, we consider how posture affects the vibrations that are transmitted to different joints in a female black widow spider, freely and naturally suspended on its own web.

Since the primary mechanical input to lyriform organs results from the bending of leg joints(3–5), it would be ideal to study joint bending directly using strain sensors. However, given the size of the legs of the black-widow spider and strain amplitudes that we expect, it is difficult to manufacture strain sensors small enough that they would not themselves disrupt the body mechanics of the spider. Laser Doppler vibrometry (LDV), however, allows contact-free measurement of movement at near picometer sensitivity(28) and we can infer joint bending from the relative motion of different leg segments under the reasonable assumption that these behave as rigid bodies at the low frequencies under consideration. We exploit the sensitivity of LDV, in combination with multi-body dynamic modelling (MBD), to study the effects of the body, its size, and posture on the tuning of leg joints in the female black-widow spider.

## Results

### Frequency segregation in the body

A vibrational input was produced by suspending a permanent magnet (of similar mass to a male spider) on the web, and by driving it with an impulsive force generated by an electromagnet. The vibrations produced in the magnet assembly were harmonic (Fig. 1A) and included a range of frequencies (Fig. 2A). Measurements were focused on the long fore-leg (leg 1) and the abdomen since these were the easiest to measure from. The resulting vibrations in the spider body were also harmonic, but different body parts vibrated at different frequencies (Fig. 1A). The vibrations observed in distal leg segments contained higher frequencies than those in the vibrations of proximal leg segments and the abdomen (Fig. 1A). Spectral analysis showed that distal segments had higher vibration velocities above ~30 Hz, than proximal segments (Fig. 2A).

**Figure 1:**
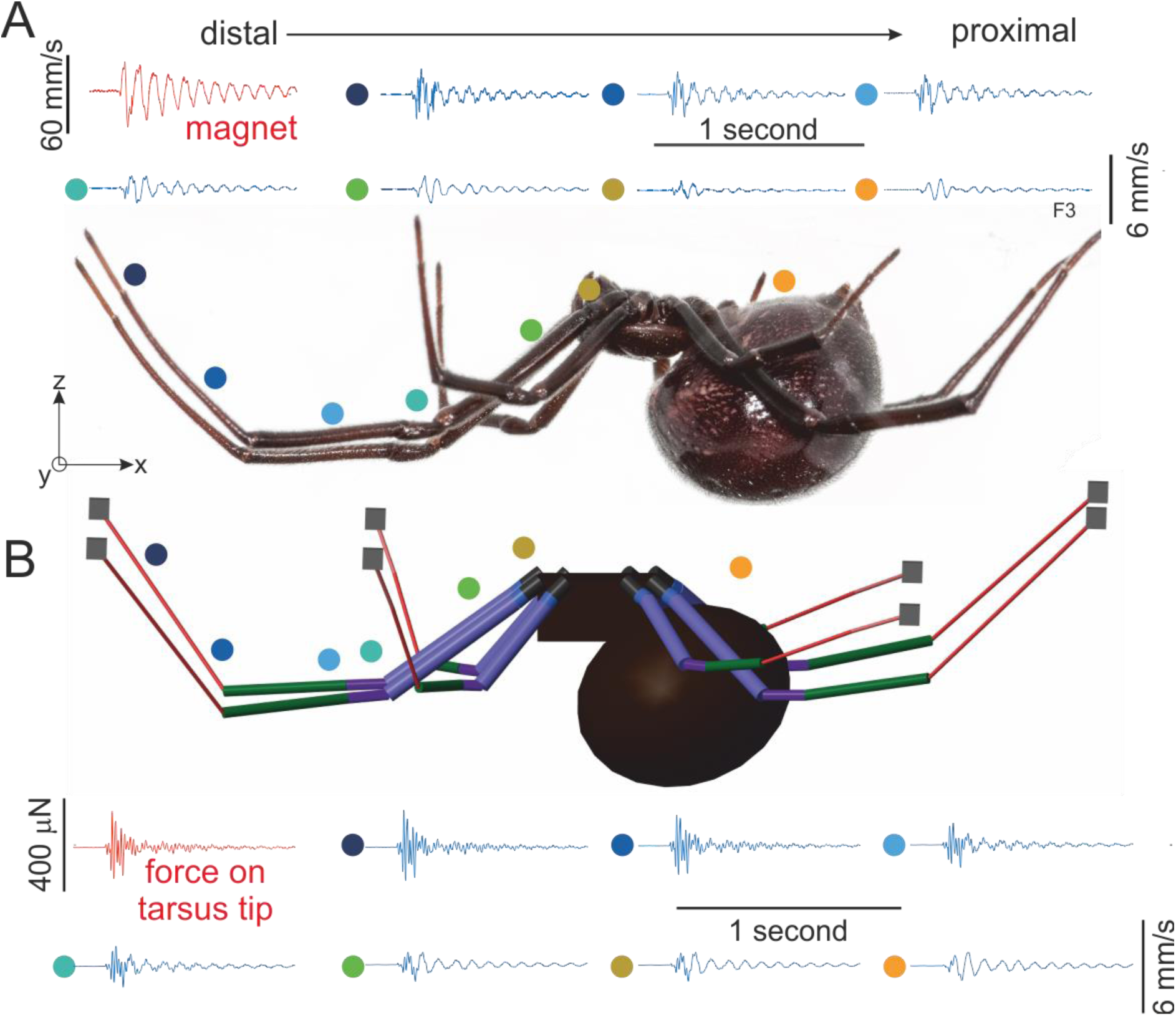
Vibration transmission through (A) a real spider’s body compared to the prediction of (B) the model spider. Waveforms of the vibration velocity of different parts of the female body are presented (position indicated by coloured circle). In the real female and in the model we observe that distal leg segment show higher frequency motion than more proximal segments and the abdomen, which move primarily at low frequencies. This shows that high frequency vibrations are dissipated at more distal joints, than are low frequency vibrations. (The motion of the permanent magnet assembly in response to an impulse force from the electromagnet is presented in red. Similarly the force applied to all leg tips in the model, which approximates the real force on the spider legs, is also shown in red.)

**Figure 2:**
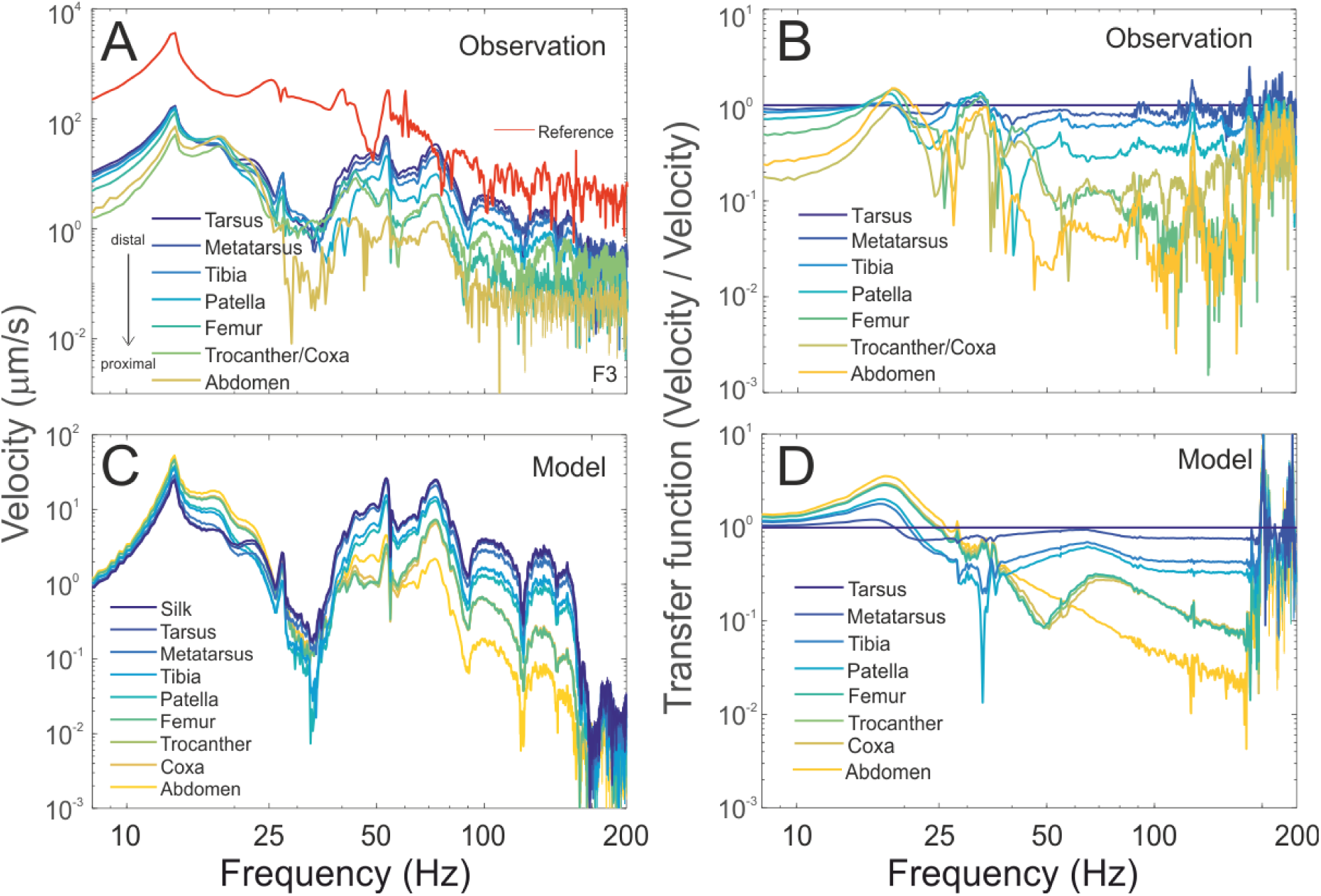
Vibration spectra allow more detailed comparison between (A) the real and (C) modelled behaviour of the vibration transmission within the spider’s body. Spectral analysis of the vibration waveforms demonstrate the segregation of vibration frequencies along the body more clearly and show that the model captures the measured behaviour of the spider adequately. In both the real and model spider, all segments show similar levels of motion below 25 Hz and above this frequency, we see decreasing motion in the proximal leg segments and abdomen. In order to estimate the dissipation of different frequencies through the body, we calculated the transfer function of the different body parts relative to the tarsus, the segment through which vibrations enter the spider body, in both (B) the real and (D) the model spider. Again we see the model captures the behaviour of the real spider well in representing the absolute levels of decay at different vibration frequencies.

A transfer function analysis was used to remove the effect of the vibration transmission through the web, and to concentrate on the biomechanics of the spider body. Vibrations are introduced into each leg of the spider at the point of contact with the web - the tip of the most distal segment, the tarsus. We therefore calculated a transfer function of the vibration of each segment with respect to that of the tarsal segment (Fig. 2B). These leg tip transfer functions allow us to estimate the decay of vibrations of different frequencies after they are introduced into the spider body and as they travel through the body. The transfer functions indicate that frequencies above ~30 Hz decay substantially as they transmit through the spider’s body. The largest decreases in high frequency vibration velocities occur at the joints between the metatarsus and the tibia, the tibia and the patella; and then between the patella and the femur (Fig. 2B). Thus, at these higher frequencies, each distal segment moves more than the next proximal segment, suggesting that the joints between these segments are being bent. A similar pattern is evident for lower frequencies, but mainly between the femur, trocanther/coxa, and the abdomen (Fig. 2B), indicating that these more proximal joints are bending at lower frequencies.

### Modelling whole-spider vibrational mechanics

We simulated vibration transmission in the spider body using multi-body dynamics (MBD) modelling. When driven by forces that mimic our experimental stimuli (see methods), the model shows similar vibrational behaviour to that of the real spider (Fig. 1B). The waveform and the levels of the vibrations are similar in corresponding body segments. Additionally, as observed in experimental measurements, the vibrations of more distal segments had higher frequencies than those of more proximal segments (Fig. 1B). Spectral analysis of the vibrations observed in the model also shows a very close match to the behaviour of the real spider, both in levels and in the overall frequency behaviour (Fig. 2A, C).

A transfer function analysis of the relative motion of leg and body segments also suggests that the model performs reasonably well at capturing the intersegmental vibrational behaviour (Fig. 2B,D). The transfer functions predict both the decay of frequencies higher than ~30 Hz as they travel through the body of the spider, and an increase in motion at low frequencies. The model, like the real female, predicts bending at the distal joints between the metatarsus, tibia, patella and the femur at high frequencies. At low frequencies, it predicts bending at more proximal joints, between the femur, trochanter/coxa and abdomen.

Our model has one important free parameter, joint stiffness, which was tuned to match the vibrational behaviour of the different segments of leg 1 (see methods, Fig. S3). Other features of the model, such as the geometry and density of body segments, are either from measurements or from the literature. This model captured the main features of the vibrational behaviour of each segment (Fig. 1, 2), including the intersegmental decay of frequencies along the leg (Fig. 2). Our results also show that the observed segregation of low and high frequencies along the leg, was not strongly dependent on the specific value of this or other parameters (Fig. S3). Thus the MBD model developed here is sufficient for predicting the segregation of vibrations of different frequencies within the long foreleg of female black widow spiders.

There were some differences between the model and the real spider, but these can be explained by the variation in locations of vibration measurement between the model and the real spider, and the subtle variation in the posture of female spiders which cannot be matched perfectly in the model (see later sections, Fig. S5).

### Full body vibrational modes

We further test the accuracy of the model, by considering its predictions over the entire spider body. The local peaks in the intersegmental transfer functions show two modes of vibration, one between 10 and 20 Hz and another between 40 and 60 Hz (Fig. 2D). The model predicts a first mode where the abdomen and proximal segments of all legs move relative to the distal segments (Fig. 3A) i.e. proximal joints undergo bending motions, whereas distal joints do not. The model also predicts a second mode where these relationships are reversed (Fig. 3B), distal segments move more and bending is experienced at the leg tips.

**Figure 3:**
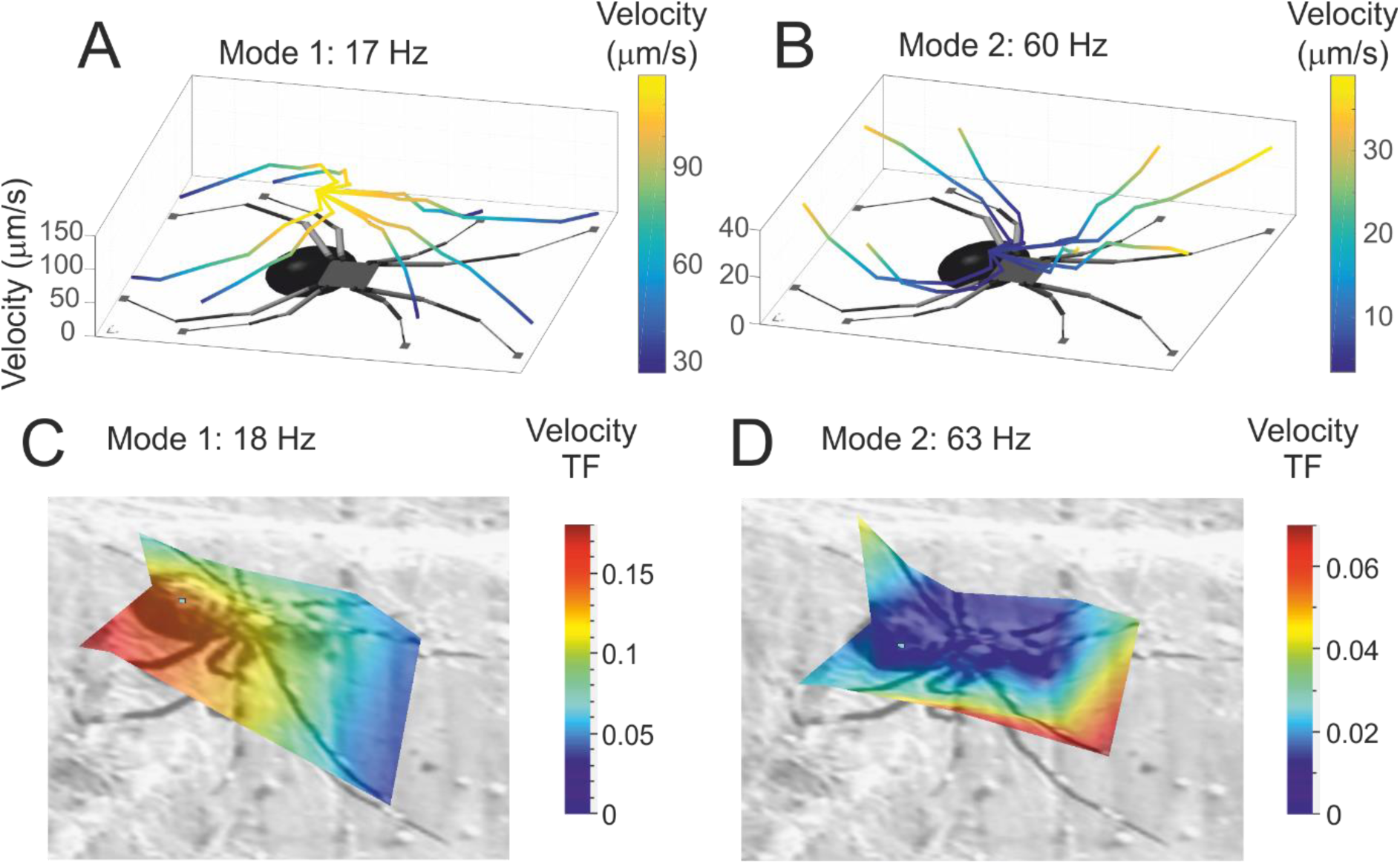
Full body vibration modes. The model predicts two main modes of vibration in the spider body. (A) The first mode of vibration is characterised by high velocity amplitudes near the abdomen, while the distal ends of the legs remain relatively motionless. (B) The second mode of motion, reverses this trend and higher velocity amplitudes are observed at the leg tips whereas the abdomen remains relatively motionless. The velocity amplitudes of the different segments of the spider body are depicted by lines whose colour is coded by their amplitude level. The first (C) and second (D) vibrational mode can also be observed in measurements from real spiders within the predicted frequency bands. (Depicted by a surface that maps the transfer function of the velocity at different points across the spider body with respect to input velocity at the vibrating magnet rather than individual leg tips.)

Scans that reconstruct the measured motion of the entire body of the spider show that these two modes can be observed in the expected frequency bands in real spiders (Fig. 3C, D; frequency of mode 1: 15.62 ± 3.66 Hz, mode 2: 55.98 ± 11.45 Hz, mean ± SD, N=10). At low frequencies, we observe motion primarily near the abdomen implying bending at proximal joints and at higher frequencies, we observe motion at the leg tips only, implying bending at the distal joints. The relative amplitudes of the two modes as observed are similar to those predicted by the model. However, in real measurements, the relative amplitudes can be highly variable (Fig. 3D). Very high amplitude motion is often observed at some leg tips. These high amplitude motions are likely the result of local variations in silk stiffness, or discontinuities in the heterogeneous structure of black widow web, which we do not incorporate in our model. Nonetheless, the model which has been tuned on the behaviour of the long foreleg is capable of capturing reasonably well the vibrational behaviour of the complete spider body and therefore can be used to predict its mechanical behaviour in other contexts.

### Effect of body size

The majority of a female’s mass is in the abdomen, which therefore represents the major inertial component governing the vibrational behaviour of the spider body. There is considerable variation in abdomen size and therefore mass among female black-widow spiders; additionally the mass of the abdomen can undergo large changes after feeding but may also increase in density (19, 29). Indeed, in black widow spiders, leg length does not vary a great deal and most of the change in spider mass can be explained by changed in abdominal size and volume [23]. We used the model to investigate the possibility that this natural variation in the spider body could affect the frequency segregation we have observed. First, we varied the size of the modelled abdomen which changed overall mass, then we changed the density causing additional changes in mass (see methods and Fig.S4).

The model predicted only minor changes in frequency segregation due to variation in abdominal size and density (Fig. 4A-D, S4). We tested these predictions by measuring frequency segregation in the bodies of real females of different masses (Fig.4C-D, S5; frequency (mode 1) = 13.47± 2.39 Hz, frequency (mode 2) = 49.14 ± 16.09 Hz, mean ± SD, N=5) and by simulating the effect of a large meal by adding a small mass (95 mg) to their abdomens externally (Fig. 4E-F, S5; frequency (mode 1) = 13.23± 5.20 Hz, frequency (mode 2) = 45.47± 13.89 Hz, mean ± SD, N=5). The vibrational behaviour of the females corroborated the predictions of the model and did not vary greatly with initial abdominal size or change after a mass was added (Fig.4G-H, S5; paired t-test for frequency (mode 1): P=0.44; paired t-test for frequency (mode 2) = 0.17, N=5).

**Figure 4:**
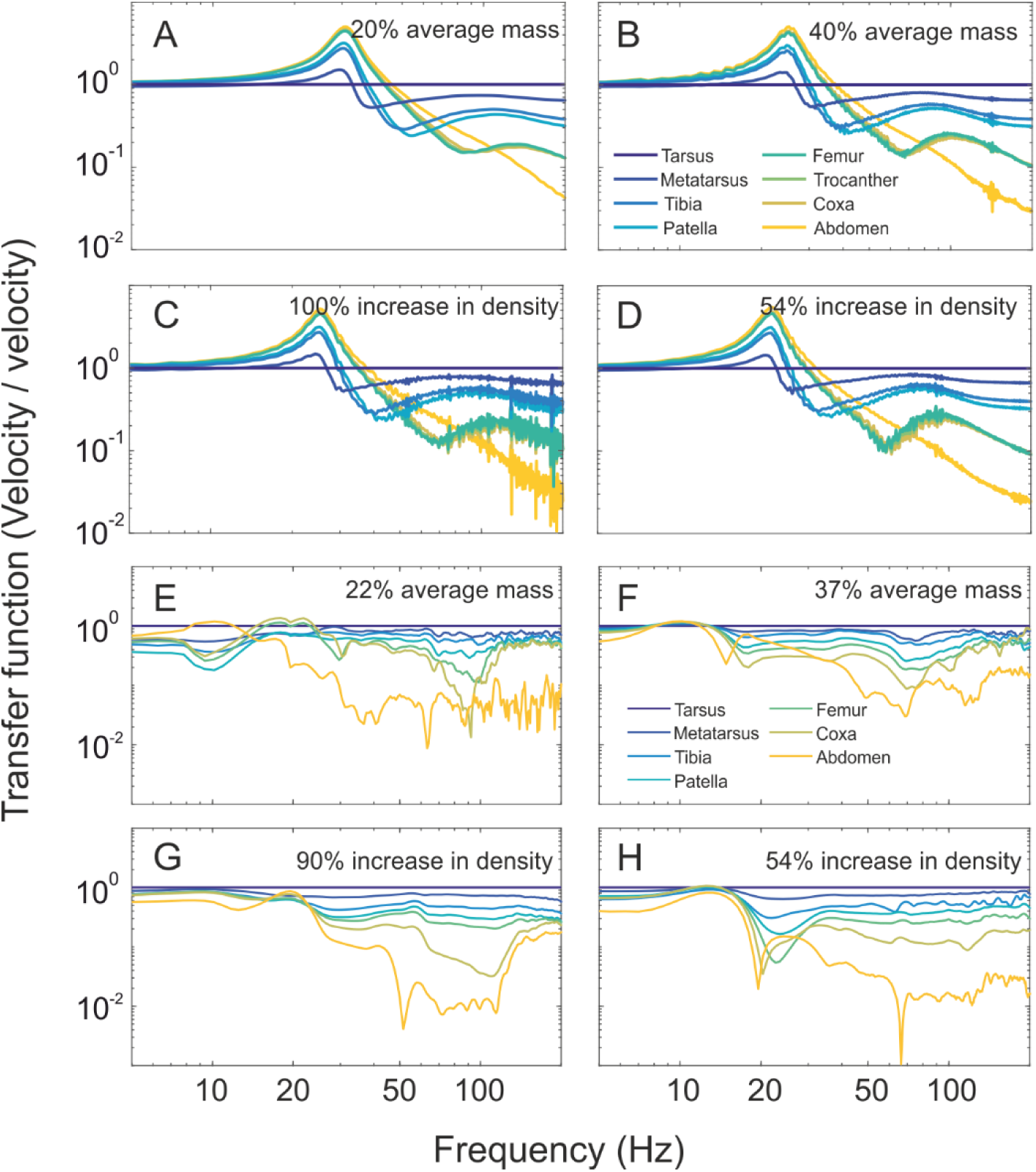
Abdominal size and density has little effect on the segregation of vibration frequencies. (A-D) The model predicts that frequency segregation within the body is not significantly changed even with abdominal mass is as low as (A) 20% and (B) 40% of the average mass (changed by changing abdominal size alone in the model). Similarly an increase in the density of the abdomen, as might be seen post a large meal also does not cause a large change in vibrational behaviour; ((C) increase of 100% and (D) increase of 54%). The predictions of the models were verified on real animals, of similar weights, (E) 22% and (F) 37% who showed similar vibrational behaviour. A small weight was then added to their abdomens to simulate a large meal. Also as predicted, neither a mass increase of (G) 90%, nor (H) 54% changed vibrational behaviour greatly.

Thus the vibrational dynamics of the black-widow spider body are robust to natural variation in body size. This might underlie their ability to correctly segregate and discriminate between incoming vibrational signals as has been suggested before by their ability to recognize courting males despite changes in feeding status(26, 30).

### Effect of posture

Posture is another obvious variation in the body of the female black-widow spider that might affect the mechanics of vibration. In general, female black-widows, like most web-dwelling spiders, are sit-and-wait predators (21, 31) and remain motionless on the web for extended periods. Nevertheless, their posture while on the web is variable, and three distinct postures are commonly observed: the “neutral posture” (legs extended, body horizontal); the “lowered-abdomen posture” (legs extended, body angled away from the web in the antero-posterior direction); and the “crouching posture” (legs retracted) (Fig. 5). Neutral posture is the most commonly observed. Lowered-abdomen is commonly observed during courtship when it indicates receptivity to a male (M. Andrade, personal communication). The crouching posture is usually adopted in the refuge part of the web and is more usual when the female is hungry or after a large vibrational disturbance on the web, (e.g. wind). In each of these postures, the abdomen, the inertial element of the spider body, is supported by legs held at different angles, which in turn gives rise to different leg spans on the web and also changes the distance from the web surface (Fig. 5). Do these subtle changes in posture change the vibratory dynamics of the spider body?

We can now use the model to investigate the effect of posture on how vibrations are transmitted and segregated through the female body. In verifying the model, we have been using transfer functions to infer the bending at different leg joints with the female in neutral posture. We have repeated these calculations for the other postures as well (Fig. S6). After running the simulations, we experimentally tested the model’s prediction in the crouch posture (Fig. S7). As with the neutral posture, there is a reasonable match between the transfer functions predicted by the model and those we observe from real animals (Fig. S7). However, since it is joint bending that drives slit compression and expansion(4, 7, 8, 32), the main quantity of interest is joint bending, which we can obtain directly only from the models. The model thus enables us to directly calculate bending velocity spectra and enables an estimation of the mechanical sensitivity of leg joints (Fig. 5).

**Figure 5:**
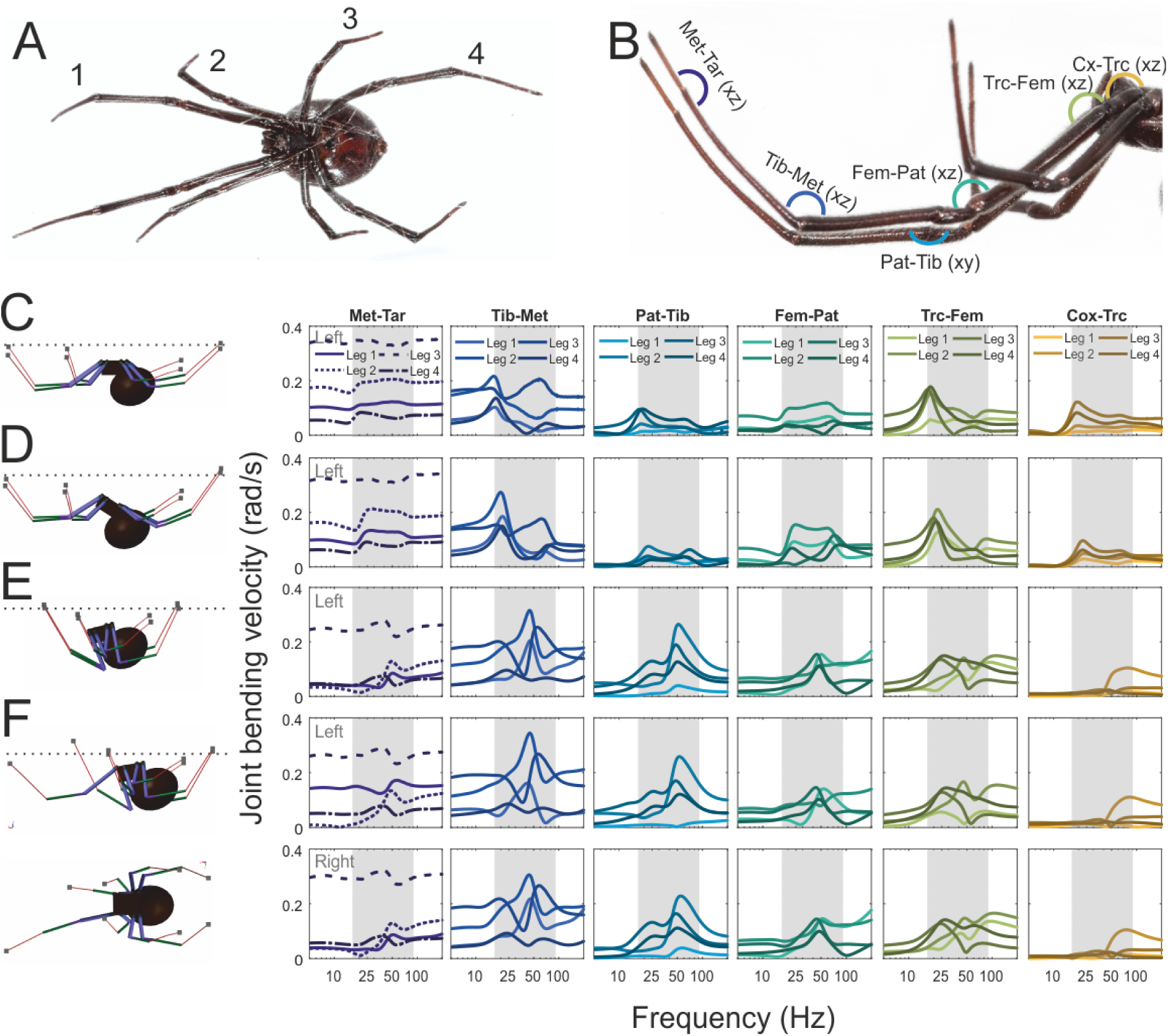
Posture affects joint tuning. Slit sensilla, including lyriform organs that surround leg joints are stimulated by joint bending. The nomenclature we use to describe legs and joints is defined here. (A) Legs are numbered from the front to the back. (B) Joints are named by indicating the two leg-segments connected by the joint in the distal to proximal order and the bending plane is indicated on the figure. (C-F) The model allows us to investigate the amplitude spectra for bending velocity at all joints in different postures (shown on the left) and spectra were plotted for four postures, (C) the neutral extended-leg posture, (D) the drop-abdomen posture adopted during courtship and (E) the crouch posture adopted usually within the refuge in the cob-web and (F) a hybrid posture with one leg extended from the crouch posture (In symmetric postures only data from the left legs are presented, and in the asymmetric posture data from both sides are presented. The side is indicated in the first subpanel. The legs are relatively symmetrical in their behaviour and each leg is represented by a progressively darker shade of the main colour. Larger differences in behaviour are observed at the first joint however, and so this joint is represented using both different lines styles and progressively darker colours. The grey shading in the spectra indicates the frequency band of male signalling (Fig. S8))

The most striking conclusion from joint sensitivity spectra is that most spider leg joints are tuned. Interestingly, this tuning is not seen in peripheral neurophysiological measurements, which in most spiders has been observed to be flat below ~100-1000 Hz(3, 4). This suggests that, although the receptors can be broadly tuned, the mechanical input to the joints shows frequency tuning. Therefore, the stimuli reaching the receptors are transformed by the body mechanics of the spider. Thus, body mechanics can play an important role during frequency discrimination in spiders.

Other insights arise from considering vibrational motion in an ‘embodied’ fashion, especially concerning the effects of posture on joint tuning. Here, we consider the pattern of sensitivity at each joint as determined by bending amplitude and frequency tuning.

#### Mechanical joint sensitivity

The tarsus-metatarsus joint is well known to be important in vibration sensing in spiders(33) since it is not bent significantly during locomotion, unlike the tibia-metatarsus or femur-patella joints. In all three postures, this joint shows very low tuning and has a nearly flat bending spectrum. However, it also experiences the largest amplitude bending velocities, and has the highest mechanical vibration sensitivity. Among the four legs, the highest sensitivity is surprisingly observed in leg 3 in all postures (Fig. 5C-F), rather than the long fore-leg (leg 1) which is often used by the female in exploratory behaviour. Nonetheless, the tarsus-metatarsus joint also tends to be the joint with the highest overall bending velocity in all legs in the neutral and lowered abdomen posture. The bending amplitude at each leg differs, which suggests that the legs may achieve a degree of range fractionation. Our results suggest that the tarsus-metatarsus joint may be specialised for detailed sensing of small amplitude web-vibrations in the context of prey capture and courtship but also for faithfully representing the full frequency range of a signal.

#### Mechanical joint tuning

All other joints are tuned, either uni- or bimodally in the frequency range of interest. As expected from studying intersegmental transfer functions (Fig. 2) and the full body vibration patterns, we find that in the neutral posture, the modal peaks occur either at ~20 Hz or ~60 Hz. The low frequency mode causes some bending at almost all joints, whereas the high frequency mode is only observed at the distal joints. The highest tuned motion occurs at the distal tibia-metatarsus joint, and the proximal trocanther-femur joint. The frequencies that the joints are tuned to in this posture form a large component of male courtship signals (Fig. 5, S8). In the lowered-abdomen posture, usually adopted during courtship, tuning remains similar, and is marked by increased mechanical sensitivity at the low frequency mode in legs 1 and 2, suggesting increased ‘attention’ to low frequency portion of male signals.

Male courtship frequency range overlaps with that of large prey in the low frequency end of the spectrum (Fig. S8,(34)) suggesting that both these postures allow females to simultaneously pay attention to prey and male signals. Some prey signals, particularly of large prey items such as crickets include frequencies even lower than this range and it is likely these signals are sensed mainly in the metatarsus-tarsus joint.

#### Crouch posture

The most dramatic changes in mechanical sensitivity spectra occur in the crouch posture. The crouch posture reduces the leg span of the spider on the web and brings the abdomen closer to the attachments on the web. This has the dual effect of severely reducing the mechanical sensitivity of most tarus-metatarsus joints and shifting joint tuning upwards, the first peak now appearing at ~50 Hz compared to ~18 Hz in the neutral posture. In the crouch posture, tuning moves to a range that overlaps the vibration frequencies usually produced by much smaller prey such as ants (Fig. 5E, S8). This suggests a possible mechanism whereby a hungry female could “tune” her peripheral vibration sensors to be more attentive to smaller rather than larger prey. More crucially, given the reduced mechanical sensitivity to all signals between 10 and 50Hz, a female in this posture is insensitive to the majority of male courtship signals, and may either ignore them or misidentify them. Thus there might be a sensory deficit only in this posture which may contribute to the observed increase in cannibalism in hungry females(29, 30). Additionally, in this posture, the female would not sense the very low frequency vibrations caused by environmental disturbances such as wind, potentially preventing receptor habituation and preserving sensitivity to other signals in other frequency ranges.

#### Hybrid postures

Given the highly irregular structure of a cobweb, females are rarely as symmetrically positioned as our models. Additionally, females in one posture are also known to partially adopt another posture, such as extending a single leg from a crouch. So next, we examined whether such a hybrid posture could combine the mechanical sensitivity of each individual state simultaneously? A single extended leg, for example, could allow vibrations to be detected more sensitively from a specific part of the web. An analysis of joint bending spectra bears out this intuition. The extended leg, which alone has a ‘neutral’ leg-extended posture with respect to the abdominal mass, regains some of the mechanical sensitivity it had in that posture (Fig. 5). The main changes are seen in the bending amplitude of the first two joints in the extended leg: the mechanical sensitivity of the tarsus-metatarsus joint increases and the frequency tuning of the tibia-metatarsus joint lowers (Fig. 5). The other legs and their joints, on the other hand, largely retain the tuning of the ‘crouch’ posture. This shows that both abdominal mass and leg position affect vibrational input to slit sensillae and that at least for the long front legs, their sensitivity may be configured somewhat independently of the rest of the body. This is consistent with the way the spiders seem to use these legs in a sensory role.

## Discussion

### Simplified body and web mechanics

In the model presented here, we have only considered the interactions between segments that are directly articulated by a joint and made the simplifying assumption that joint stiffness is uniform throughout the body. While we have tested the sensitivity of the model to some of these parameters and found it to be relatively robust (Fig. S3, S4), we know that this does not fully describe the joints of spiders whose stiffness can change with posture(1) and whose visco-elastic properties affect the transmission of vibrations through the leg(35). Additionally, there might be joint configurations, other than the revolute joint, that may be more appropriate to capture the more detailed temporal dynamics and frequency segregation of the spider leg(32). Besides the details of joint behaviour, there may be other intersegmental interactions that occur due to contact between disconnected body parts which we have not considered.

Spiders in general (17) and black-widow spiders, in particular,(19, 22) are known to change the structure of their webs and silk in response to their condition and sexual status. This has been suggested to be a form of extended perception rather than the embodied perception that we have presented in our work(17, 23). In effect, we have taken a simplified approach to studying spider body mechanics, and have used internal transfer functions to isolate the effect of the body from that of the web. Despite the simplifications, the model we have developed captures reasonably well the overall vibrational behaviour observed from real spiders. The model also predicts significant postural effects, and these mechanical features, along with the web structure, may permit the spider additional control over perceptual processes which are likely to have been underestimated by our simplified approach.

### Multifunctional sensors

Many organisms control proprioceptive feedback via descending neuronal systems which typically refine the output of locomotor behaviour(36). In our work here, we attempt to close the sensorimotor loop and show how sensory input can be modulated through locomotor and behavioural mechanisms. Our results demonstrate how both the levels and tuning of the mechanical inputs to spider lyriform organs can be reconfigured simply by changing posture. Reconfigurable mechanical tuning may reflect a need for slit receptors (i.e. lyriform organs) to precondition the stimulus for the nervous system. Control over the mechanical input to the nervous system would allow them to be multifunctional; i.e. function in different behavioural contexts, such as locomotion as well as prey and mate perception(15). Each of these contexts requires different amplitude and frequency sensitivities. Locomotion generates larger leg bending motions than web vibrations would and at much lower frequencies(4). It is very likely that lyriform organs are activated during locomotion, and may be used to provide feedback and proprioceptive control as they are in the large wandering spider, *Cupiennius salei* lyriform organs (1, 4, 37). Similarly, prey caught in the web are likely to make large struggling movements and produce high amplitude vibrations(26) at a range of frequencies, which likely depend on prey size (Fig. S8). The lowest amplitude vibrations are likely to be those made by males(26). Additionally, male signals may contain frequency dependent information crucial to determining male identity and quality. Our results suggest that female posture affects both joint bending amplitude and frequency tuning, and this property may be deployed by females in different behavioural contexts. It would be interesting in the future to examine the mechanics and peripheral tuning of the posture adopted by females in each of these contexts in greater detail.

### Sensory complexity and intentionality

Using a dynamic systems approach it has been suggested that each posture in an animal’s repertoire is a state of neuro-mechanical equilibrium, and that animals can transition from one posture to another depending on the sensory feedback they receive(15). Different postures may also be viewed as points from which locomotory programs may be initiated, with some being more likely than others. An interesting possibility from this kind of description is that static postures can be thought of as equilibria. Some equilibria will be closer to each other than others in “posture space”. In the black widow system, for instance, joint tuning suggests that the lowered-abdomen posture allows the female to be more sensitive to male signals. Therefore the dynamics of bodies may have evolved such that they can transition preferentially between relevant postures i.e. equilibria(15). For instance, the lowered-abdomen posture might both allow the female to perceive the male better, but also allow her to transition into the mating posture. Similarly, while the crouch posture allows the female to perceive even small prey items, it might also allow her to transition into an attacking posture more easily.

Additionally, the transitional motions between equilibria can also be mapped as existing in joint angle space. Since transitional postures will be small continuous changes in joint positions, which define their position in this multi-dimensional space, they are expected to lie on a continuous but bounded surface within this space, i.e. on a manifold. In kinematics, postural manifolds are routinely used in machine vision to identify postures visualised from different angles, and in robotics, to plan the actions in a multi-limbed and jointed body(38–40). Recently some evidence has been found for neural manifolds controlling motion in biological systems(41). Our data suggests that posture can profoundly affect the sensory system of an animal, and that transitional states between postures take on transitional sensory properties. Thus, just as one can posit a postural manifold, one can consider a sensory manifold for in a spider’s body. As our data suggests, the body and posture behaves as a sensor which has non-static properties, and pre-conditions the nature of the vibrational input to the many joints in the spider body. Therefore, interpreting the amplitude and spectral information in incoming sensory data should pose a significant challenge and should require feedback and mapping between the postural and sensory manifolds. The more complex the mechanics of the physical body, in terms of number of joints and sensors producing sensory inputs and their possible sensory states, the more complex the task of interpreting incoming sensory data is likely to be. This tight loop of interaction between perception and behaviour required by such mapping may well be what explains the surprising cognitive complexity observed in spiders(17, 42, 43).

## Methods

### Female body dynamics: vibrometry

Adult female *Latrodectus hesperus* were released onto custom-made wooden frames on which they made webs. Wooden frames were made up of a base (length × width × height cm^3^ ~ 12 × 12 × 1.5) with four posts (diameter × height = 0.8 × 11 cm) mounted at each corner (Fig. S1)). All females were given at least two weeks to construct webs. Vibrometry experiments were carried out on 10 unrestrained females in their naturally adopted posture on these webs. Females were briefly removed from the webs, anaesthetized with CO2, and the ventral surface of all leg-segments, cephalothorax, and abdomen were spot-painted with white nail polish to enhance reflectivity. Care was taken to avoid painting over the joints. Additionally, we placed a few reflective glass beads (45-63 µm bead diameter; Polytec Inc., P-RETRO Retroreflective Glass Beads) to further enhance reflectivity. Females were released back on to their webs and measurements were made only after females had recovered from anaesthesia.

To excite vibrations in the web, we used the attractive and repulsive force that can be generated by an electromagnet acting against a permanent magnet. The permanent magnet was a small neodymium disc magnet (0.25 mm diameter x 0.25 mm thickness, 14 mg mass) and was mounted on small strip of velcro (hooks side). This assembly (total mass 20 mg) was suspended on the black widow’s web at a randomly selected position. The weight and size of this assembly resembled that of a male black widow spider, or small prey, producing the same local tension in the web. An electromagnet powered by a power amplifier (Brüel and Kjær, Model Type 2718) was used to drive this assembly. The electromagnet was held 1-3 cms above the web (Fig. S1) and a 25 ms pulse was used to attract and then release the permanent magnet. The motion of the permanent magnet assembly was monitored using a single point fibre-optic vibrometer (Polytec OFV 511 with 3001 vibrometer controller) positioned above the web and at a 45° angle to the xy plane (Fig. S1). The signal was amplified until a peak velocity of ~20 mm/s was reached.

Vibration velocities (z direction) of different parts of the *Latrodectus hesperus* female were measured using a scanning laser Dopper vibrometer (Polytec PSV 400, OFV 505 scanning head) by bouncing the laser off front-silvered mirrors positioned above the web at a 45° angle to the xy plane (Fig. S1). We monitor vibration velocities rather than displacement because it enables us to consider the kinetic energy dissipation (½mv^2^) within the system, whether translational or rotational. We also monitored and retained data on signal quality along with the vibration velocity and only high quality signals were used for analysis (Fig. S2). Measurements were made in the time domain, i.e. and data were collected through the Polytec Vibrometer software (Version 9.1.1) and digitized through a National Instruments card (PCI 6110) at a sampling rate of 25.6 kHz. A total of 2.56 s of data averaged over 10 measurements was collected per scan point.

Scans over the complete body were made for 10 females in order to estimate the frequencies of the two modes. Modes were identified by mapping the transfer function between body and magnet vibrations over the spider body. Peaks in the transfer functions were identified and modes determined from the spatial map. To conduct studies on the effect of size, 5 female with below average body mass were chosen (Fig. S5) and their vibrational behaviour was tested either by conducting scans over the complete body or along the leg 1 and abdomen transect depicted in figure 1. After this a small metal washer of 95 mg was glued to the abdomen of each female, simulating the effect of a large meal. The vibrational behaviour of the female was then retested under these conditions. By beginning with small females, we did not exceed the average mass of a black widow female, and thus did place an unnatural strain on the web. By beginning with females of a range of weights, we could also test a range of percent changes in female density.

### Female body dynamics: modelling

A simulation of transmission of vibrations within the spider body was developed using a multi-body dynamics approach with the Simscape Multibody package (version 4.9) in Matlab (version R2016b). This approach treats the spider’s body segments as rigid solids connected by joints which allow relative motion between the connected segments. Spider legs are partial hydrostats. Leg extensions are achieved by increasing internal pressure, however, retractions or indeed the prevention of extension at individual joints which allows different postures to be achieved, are driven by muscles. A simple rotational-spring-mass-dashpot model such as used by an MBD approach has been found to be sufficient to model spider leg joints based on previous measurements (32, 44). In this system, the main rotational stiffness and damping is provided by the muscle, and internal extension forces are provided by pressure (32, 44). Each of the leg segments (distal to proximal: tarsus, meta-tarsus, tibia, patella, femur, trocanther, and coxa), the cephalothorax and abdomen were treated as rigid bodies. The size and position of each rigid-body in different postures was approximated from photographs and the density of spider cuticle is taken to be 1060 kg/m^3^(44). The abdominal mass of the model female was 464 mg which compares with the average mass of adult females(29).

Based on the behaviour of real spider legs, and simplifying assumptions, all inter-segmental joints within the leg are modelled as revolute joints, all rotate in the xz plane except the patella tibia joint which rotates in the xy plane(32, 45). The coxa-cephalothorax joint, and the joint between the cephalothorax and abdomen are all modelled as a ball and socket joints(32, 45). For a first approximation, and in order to avoid overfitting the model, we assume that all joints have the same stiffness, but that frictional damping would depend on the surface area of the joint. We use the joint stiffness and damping from the *Phrixotrichus roseus*(44) as a starting point and vary these two parameters to fit data from vibrometry (Fig. 2, S3). We find that implementing a joint stiffness of 30e-5 N*m/rad and damping of 1e-6 x segment radius N*m/(rad/s) replicates the mechanical behaviour of the black widow spider female and is within the range suggested by data from other spiders (1, 44). An analysis of sensitivity to these parameter choices was made and the model outputs were found to be robust to the expected range of variation in these parameters (Fig. S3, S4).

To achieve the different postures that females adopt in the model, we define the observed leg joint angles in those postures as equilibrium positions, and allow the model to reach equilibrium before testing its vibrational behaviour. Such postural equilibria have been posited in *Aplysia* and there is some evidence to suggest that these are neuromechanical in nature and achieved by coalitions of muscles(15). Spider leg joints are known to have limited ranges of motion, and to show some deflection dependent stiffness. This deflection dependent stiffness is largely with lateral deflection(46), against the bend of the joint. In axial rotation that we study here, joints are very linear(46) and only change at very large deflections, as large as 80° (1). This change is likely brought about through the recruitment of other tissues as additional spring elements, such as cuticle, or as segments contact each other. In our model, some change in stiffness is possible in such as extreme postures as the crouch position, but in the interest of parsimony, we have not included this change in the current model. Similarly, we implement a module that captures the stiffness and damping forces(47) that might arise during contact between the hind legs and abdomen, however, these are also not in play in our simulations.

Finally, while we do not explicitly model the spider web, the local attachment of the spider to web-silk is modelled as a ball and socket joint with no stiffness or damping which we believe is reasonable since this joint is actually a claw with which the spider is suspended from the silk, and which is actively controlled and repositioned by the female. We capture the local elastic behaviour of the silk itself by treating it as a prismatic joint which allows motion only in the z direction. The stiffness and damping of the web was taken from measurements of 5 black widow webs. The permanent magnet assembly was excited as described before, and an FFT of the displacement response was calculated at a frequency resolution of 0.4 Hz using a rectangular window. We tested 5 webs at 5 positions each. We fitted a simple harmonic oscillator model to the response of the web and calculated the local spring constant and damping of the web in the z direction. The webs had a mean spring constant of 0.31 ± 0.29 N/m (mean + SE, n=5 webs) and a mean damping constant of 0.06 ± 0.01 N/(m/s) (mean + SE, n=5 webs). These measures of the spring constant are commensurate with the measured Young’s moduli of *Latrodectus* silk(48).

To simulate the forces experienced by the spider on the web, we apply forces only to the tips of the model spider legs. Two types of forces are used to excite the model to study its frequency response. The first is a force similar to that experienced by the real spider in the experiment (Figs. 1, 2). We take the derivative of the velocity waveform measured from the tarsus, to calculate its acceleration which will be proportional to the local force. We multiply this waveform (5) with a constant calibrated to the output velocity amplitude and apply it to leg tips in the spider model. Both the waveforms and spectra of the model response are then compared to the measured behaviour of a real spider (Figs. 1, 2). To study the complete vibration response of the spiders body from the model, a more idealised force waveform is used, a force impulse of duration 0.1 ms and amplitude 5 N.

### Male and prey signals on web

We compared previously recorded courtship vibrations of male *L. hesperus*, to vibrations generated by struggling prey on the web (using prey items that readily elicit attach by female widows). Specifically, we used male abdominal tremulation signals produced near the female, as these signals were the most distinct and conspicuous of their repertoire. Data were collected from 16 males, 5 crickets (*Acheta domesticus*), 5 black ants (collected on pavements near the University of Toronto Scarborough; species unknown) and 5 mealworms (*Tenebrio molitor*) released on black widow webs. All prey was released ~7 cm away from the female. Web vibrations were measured near the females’ refuge in order to estimate the signal reaching the female after transmission through the web, rather than the local vibrations produced near the vibration source. Vibration data were collected using a laser Doppler vibrometer (LDV; Polytec PDV 100; 20 mm/s peak measurement range; 0.5 Hz - 22.0 kHz frequency range; 24 bits). Vibration signals were recorded from a tiny reflective tape (~1mm^2^) that was placed carefully on web-silk near the entrance of females’ refuge. The reflective tape use used to enhance reflectivity. All vibration signals were recorded and stored onto a data recorder (Sound Devices 722; 16 bits; 48.0 kHz audio sampling rate). For each of the prey item 1s of data were analysed and for each male, we analysed at least 5 sections of data of 1 second each. An estimate of the power spectrum density was calculated using Welch’s method at a resolution of 2 Hz for every section of data. For the males, the average power spectrum for each male is presented along with the population average (Fig. S8 A) and for the three prey species, data for each individual is presented with the population average (Fig. S8 B, C, & D).

## Authors’ contributions

NM and SS collected data and analysed results; NM carried out the modelling and all authors contributed to the design of the experiments and the writing of the manuscript.

## Ethics

The work reported here meets all the legal requirements and all institutional guidelines for the use of invertebrates in the country in which the work was carried out.

## Acknowledgements

This research was funded by NSERC Discovery grants to AM. NM acknowledges the support of a Wissenschaftskolleg zu Berlin fellowship in developing expertise required to use the modelling tools used here.

## Supplementary materials

**Figure S1:**
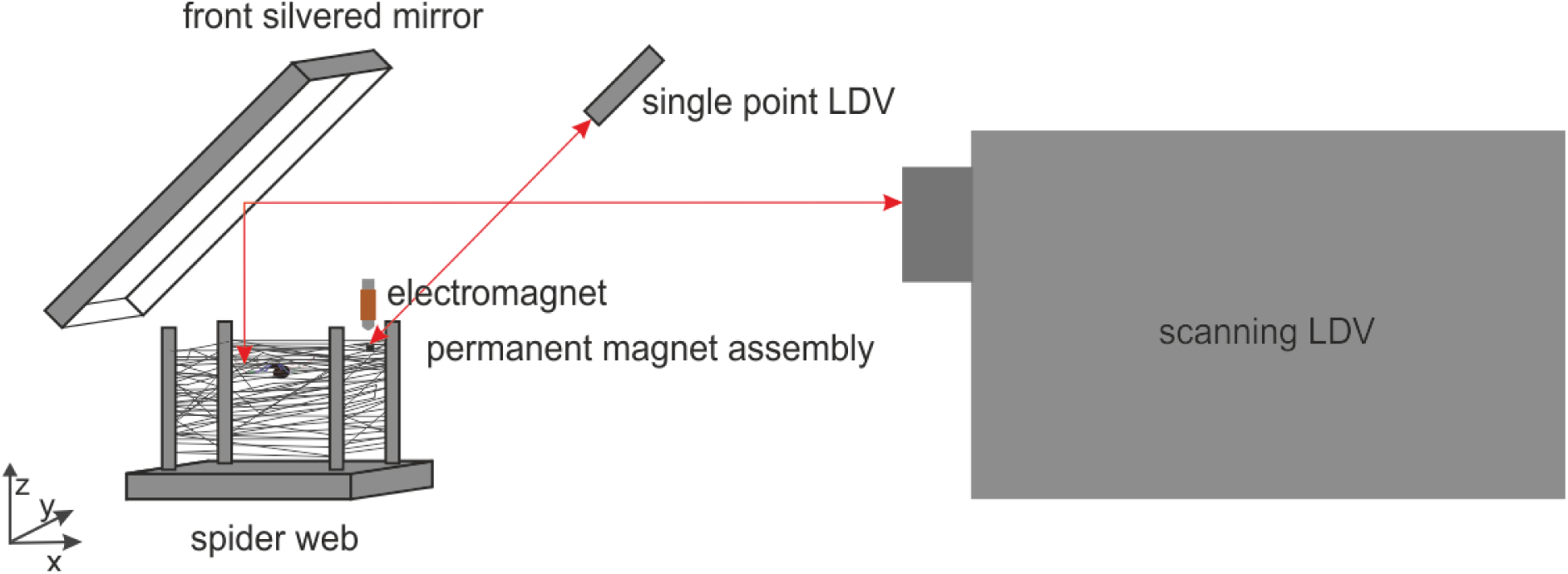
A schematic of the experimental setup. The drawing is not to scale.

**Figure S2:**
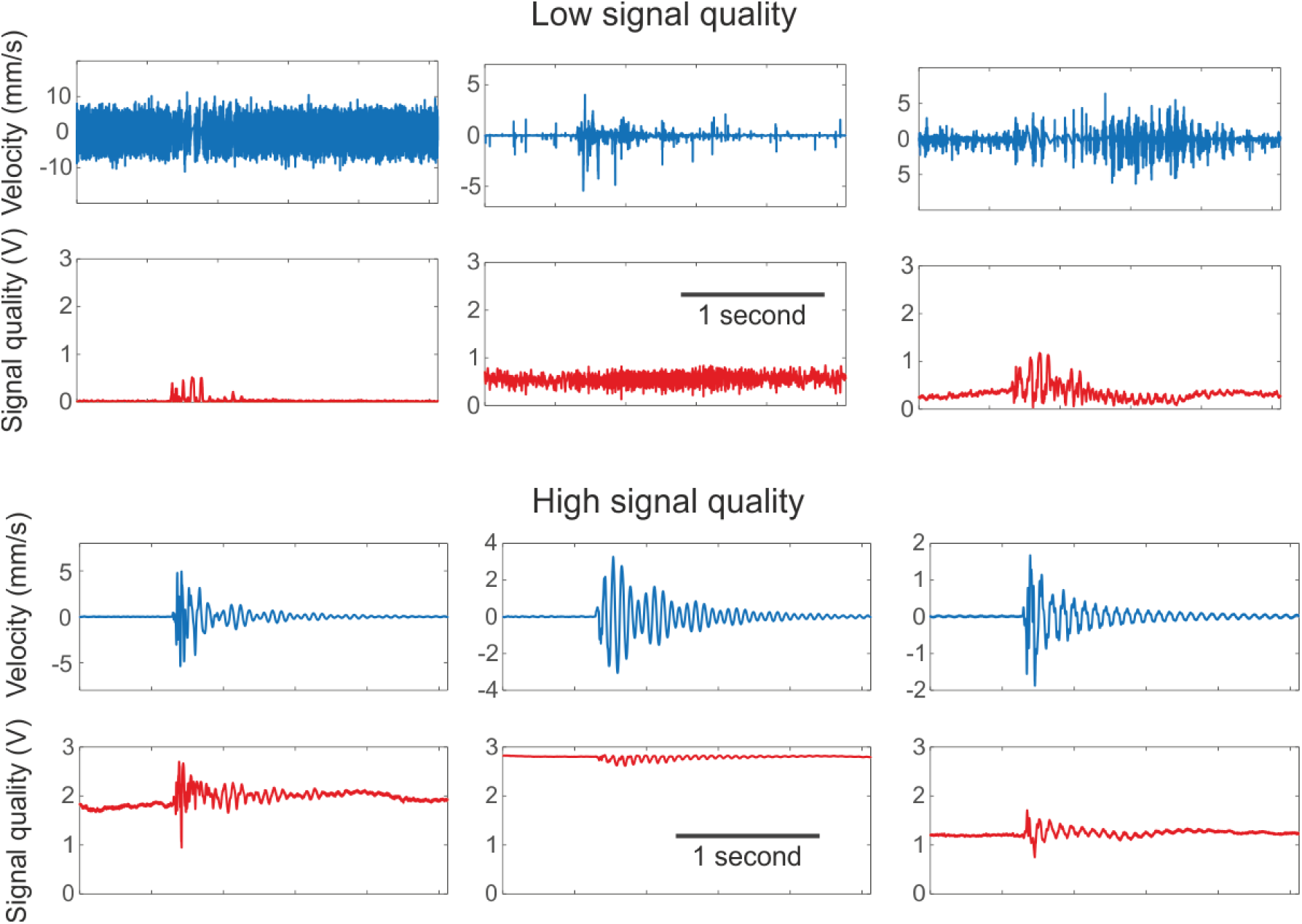
Signal quality during vibrometry. Both spider cuticle and web silk have low reflectivity and it is crucial to monitor signal quality during vibrometry in order to get reliable results. For this reason, we made our measurements in the time rather than in the FFT domain. We maintained an averaged waveform and inspected the waveform for smoothness and to detect noisy transient signals. It is worth noting that where the reflectance of the vibrating surface is low, the noise floor increases considerably (see 1st panel) and the noise may in fact incorrectly show ‘vibration’ levels higher than the real signals. If only an FFT analysis is carried out, without an inspection of the raw waveforms, the spectra would simply be of the noise, which would not be classified as such without a coherence analysis. To ensure the reliability of our signals, we monitored the signal quality measured by the scanning LDV at all times using the inbuilt quality channel which reflects the level of incoming reflected laser signal. We found that when the quality signal remained above 1V on average, the velocity outputs had smooth and continuous waveforms and did not show noisy transients. Signals below this quality level were deemed unreliable and not used in analysis.

**Figure S3:**
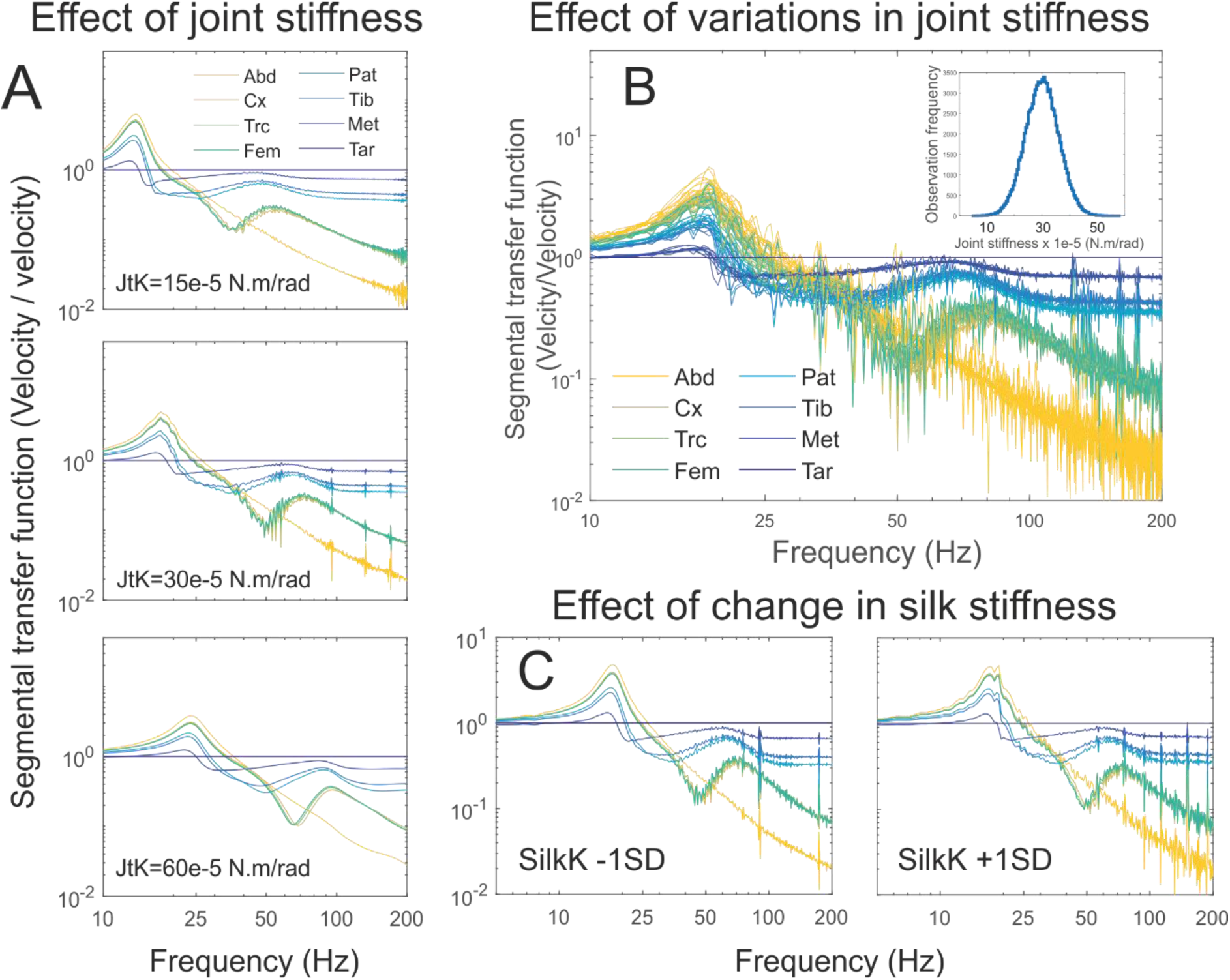
Model stiffness sensitivity analyses. The model was tested for its sensitivity to the stiffness parameters used. (A) We have used the same stiffness for all model joints and the main effect of the decreasing or increasing the joint stiffness (JtK) is a frequency shift in the transfer function spectra pattern. We test the model at stiffnesses ranging from half the best-fit stiffness to twice the best-fit stiffness. Lower stiffnesses lead to a lower peak in the transfer function and higher stiffnesses lead to a higher peak. (B) We also tested the effect of variation in joint stiffness and the results of 10 runs are plotted here. We found that joint stiffness variation had relatively little effect on the overall pattern of the transfer-function spectra. The stiffness of each joint in the model was selected from a distribution with a mean of 30e-5 N.m/rad and a standard deviation of 15% i.e. 4.5e-5 N.m/rad (inset). (C) We also tested the effect of variation of silk stiffness on transfer function spectra. The spring constant of the all attachments to web-silk (SilkK) were decreased and increased by 1 standard deviation and we found the effects on intersegmental transfer functions to be negligible. (Abd: abdomen; Cx: coxa; Trc: trocanther; Fem: femur; Pat: patella; Tib: tibia; Met: metatarsus; Tar: tarsus).

**Figure S4.**
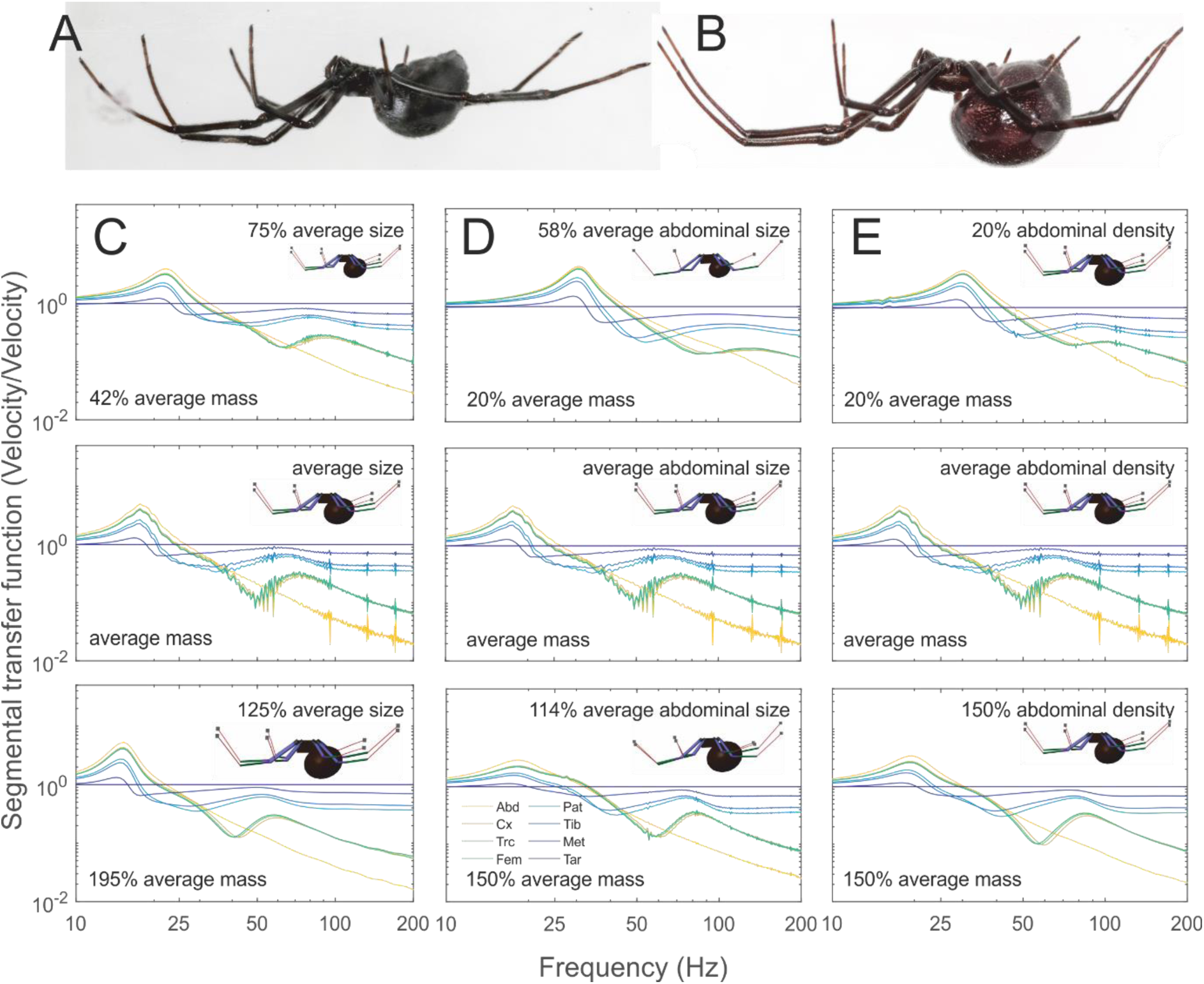
Body size sensitivity analyses. (A, B) The size and mass of a female spider can and does change in many ways. (C) One possibility is that different spiders are simply allometrically smaller and larger than each other, i.e. their spatial dimensions change by the same factor. We tested the effect of allometric size and found that larger spiders had lower peaks in their transfer function spectra but the change was small despite large changes in size and effective mass. In real black widow females however, leg length does not change a great deal (<1%) even when large changes in mass (50-200% average mass) are observed [23], implying that the change in mass occurs either through a change in abdominal size or density. (D) We varied the size of the abdomen in the model to mimic the body size variation observed in real females and found effects similar to that described before, larger females had lower frequency peaks than smaller females, however, the absolute change in frequency segregation in the spider body was small. (E) We also tested females with abdomens of different densities and observed similar model behaviour. This shows that variations in other parameters such as mean joint stiffness or changes caused by posture exceeded any variation that may be caused by size. (Abd: abdomen; Cx: coxa; Trc: trocanther; Fem: femur; Pat: patella; Tib: tibia; Met: metatarsus; Tar: tarsus).

**Figure S5:**
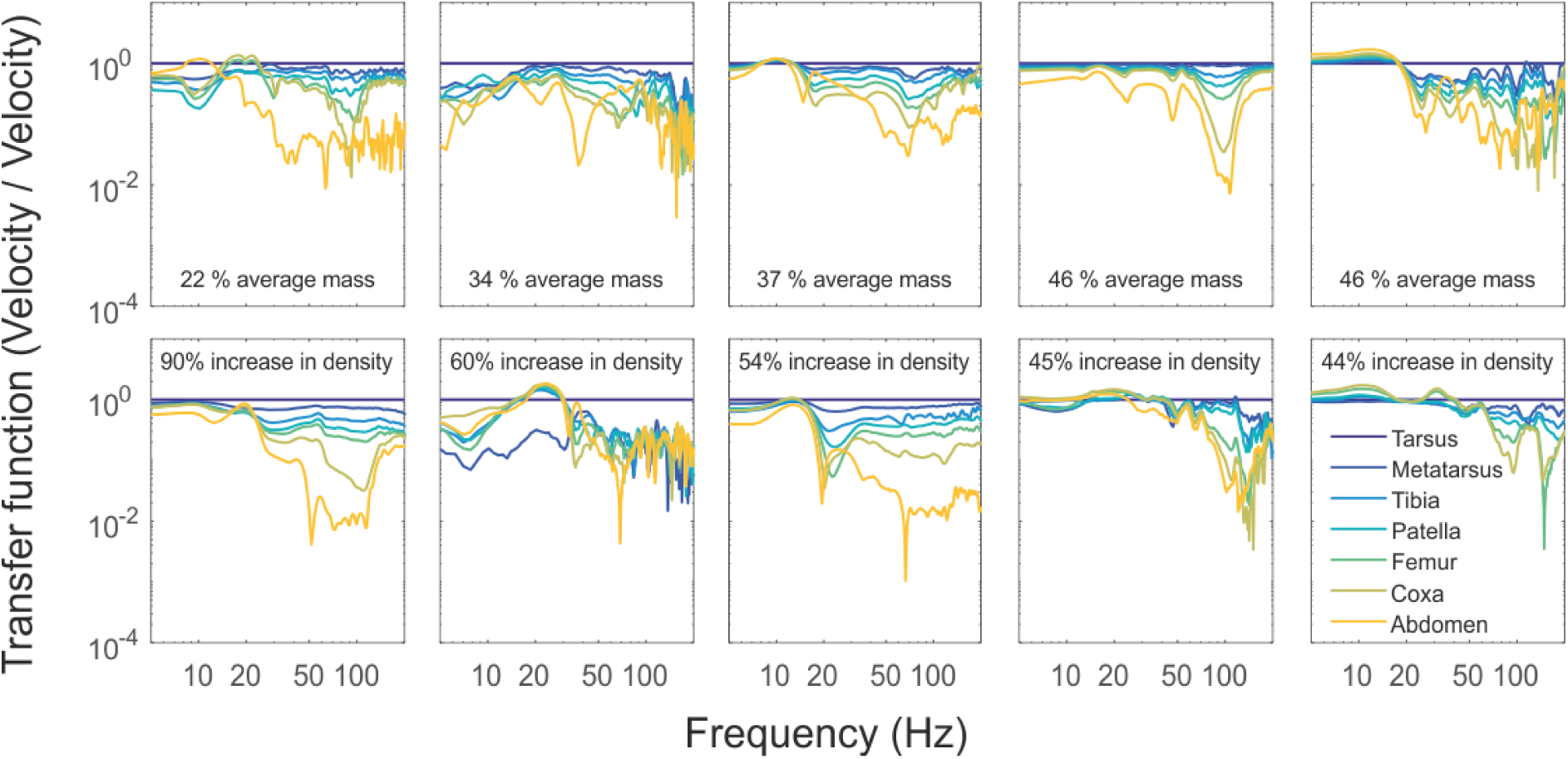
Body size variation has little effect on frequency segregation in the spider body. The model predicts that, despite large changes in abdomen mass, little change will be observed in the frequency segregation behaviour of the spider body, as indicated by a transfer function between the tarsus and other segments. We tested females of a range of body sizes and found that the peak transfer frequency variations did not change systematically with size. The first row of transfer functions are from 5 different females of increasing mass. The second row of transfer functions are from the same females after a large mass (95 mg) was added to their abdomen. Frequency segregation behaviour does not change systematically in either row of transfer functions. A lack of systemic change in frequency segregation indicates that rather than mass, a different parameter explains the variations we observe in vibrational behaviour. We believe that both variations in average joint stiffness, and changes caused by posture exceed any change that may be caused by size.

**Figure S6.**
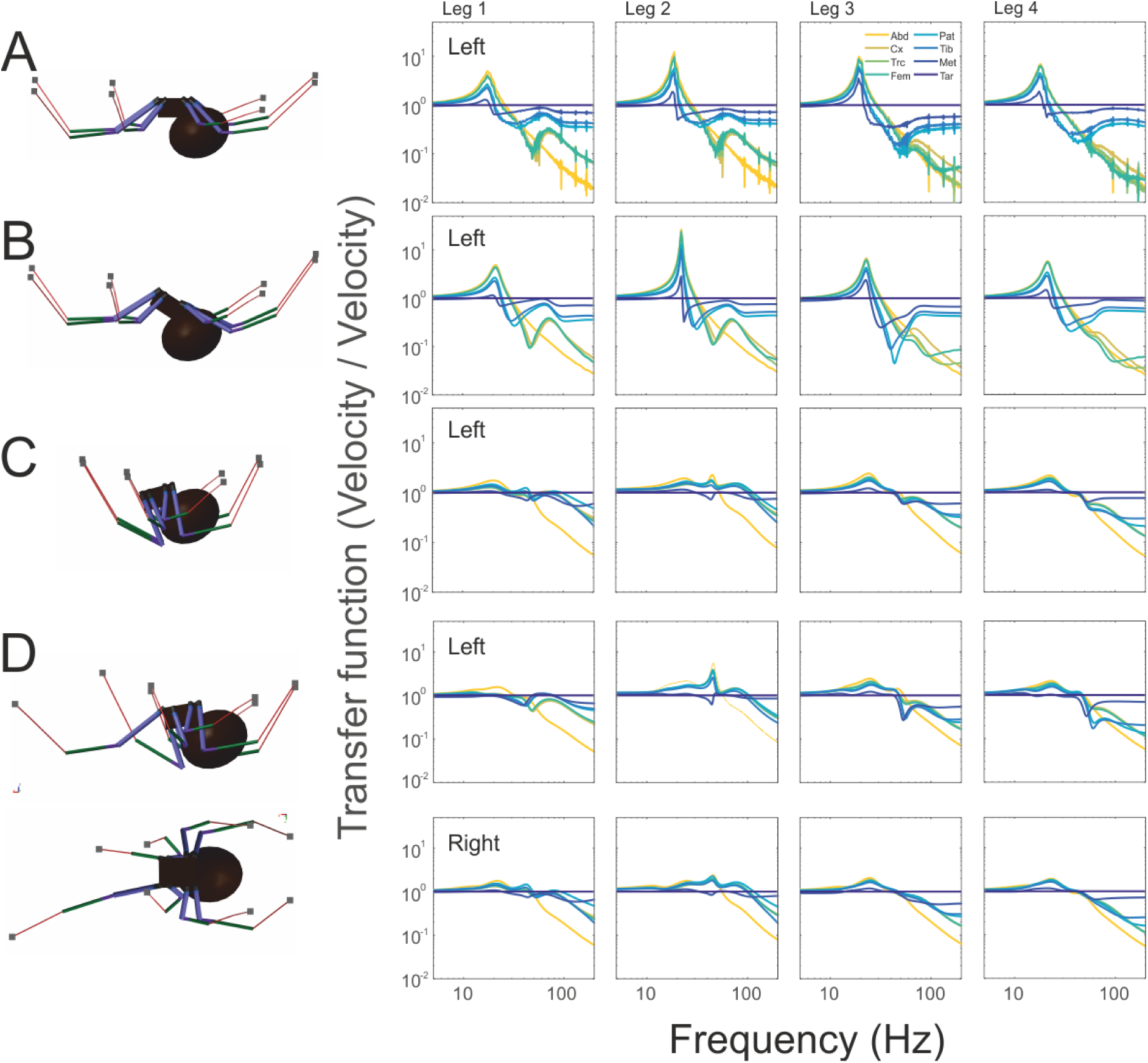
Posture has a strong effect on frequency segregation in the spider body. The segregation of frequencies in the spider body is different in the four postures investigated (A) the normal leg-extended posture, (B) the lowered-abdomen posture, (C) the crouch and (D) the hybrid crouch posture with a single extended foreleg. The main difference between the first two postures is an increased motion in proximal segments to low frequency vibrations. The main response in the crouch posture is a much lowered response to low frequencies across all legs. Differences between the hybrid and the complete crouch position can be observed but their effect is clearest in the joint bending spectra. Some of the transfer functions measured from real females, resemble the hybrid transfer functions more closely than those of the symmetric leg-extended posture which seems reasonable considering that completely symmetrical postures would be difficult to achieve on the web which provides a very sparse and irregular substrate. (Abd: abdomen; Cx: coxa; Trc: trocanther; Fem: femur; Pat: patella; Tib: tibia; Met: metatarsus; Tar: tarsus).

**Figure S7:**
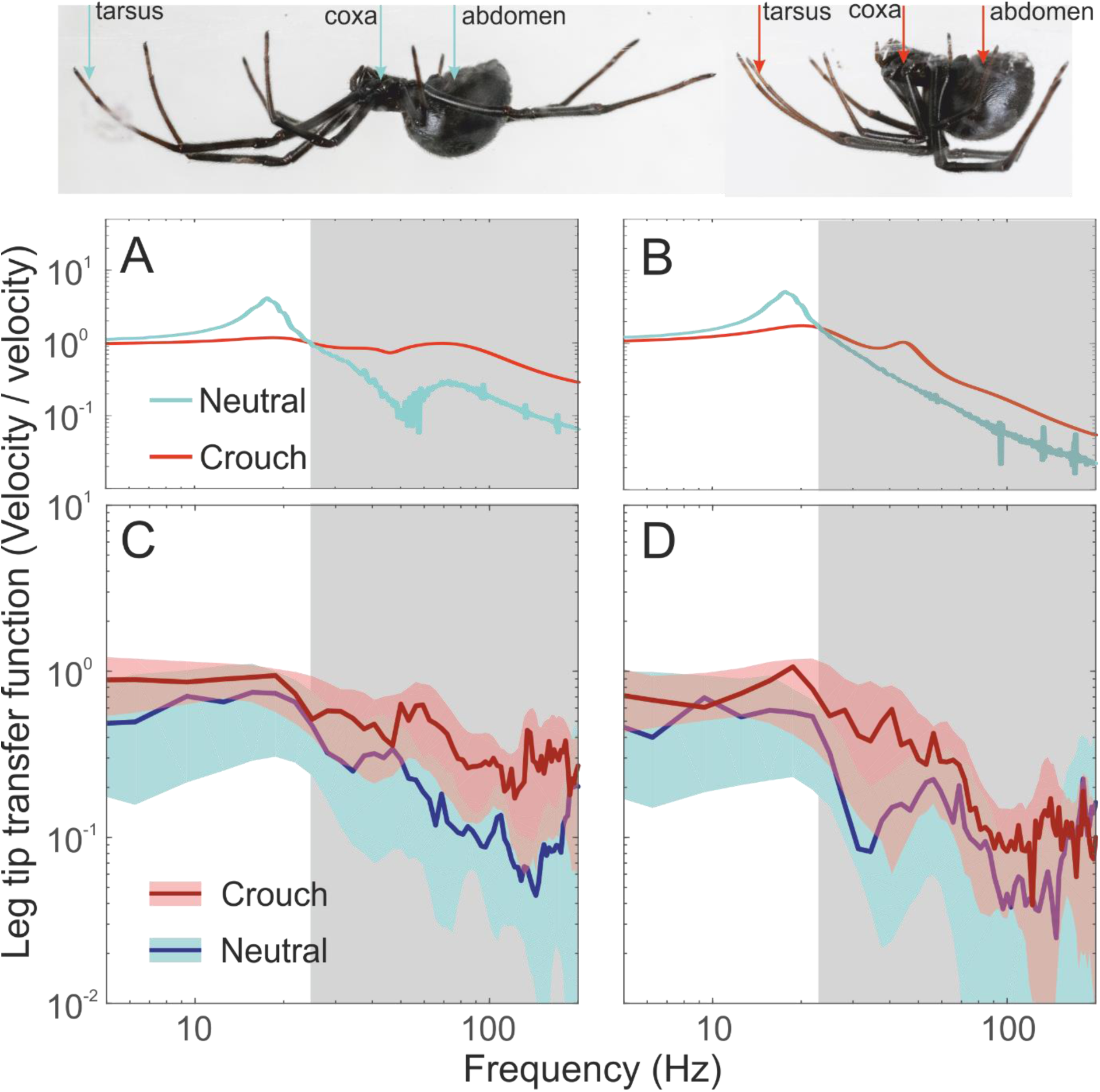
Model predictions were tested by measuring from six females in both the neutral and crouched posture. In the crouched posture, the female withdraws her legs and tucks them close to the body. In this posture, it is not possible to reliably measure from the legs. However, we can measure from the leg tips, coxa and abdomen. We compare the predictions of the model in terms of the transfer functions between the leg tips and the coxa and abdomen with real measurements. (A) The model predicts that at frequencies above ~27 Hz, the motion at the coxa will have decayed considerably more in the neutral posture compared to the crouched posture. (B) The effect at the abdomen is similar but of smaller magnitude. We find that the measured behaviour of the (C) coxa and (D) the abdomen above ~27 Hz are similar to that predicted by the model. The lines indicate the means from 6 females and the shaded region depicts the standard deviation around the mean. We make a statistical comparison by summing the transfer function in frequency range 27-200 Hz and find the measured transfer function ratio to be higher in the crouch posture (coxa: 1800±979, N=6; abdomen=123±890, N=6) compared to the neutral posture (coxa: 829±539, N=6; abdomen=720±521, N=6; paired t-test: coxa: P=0.065; abdomen: P=0.066).

**Figure S8:**
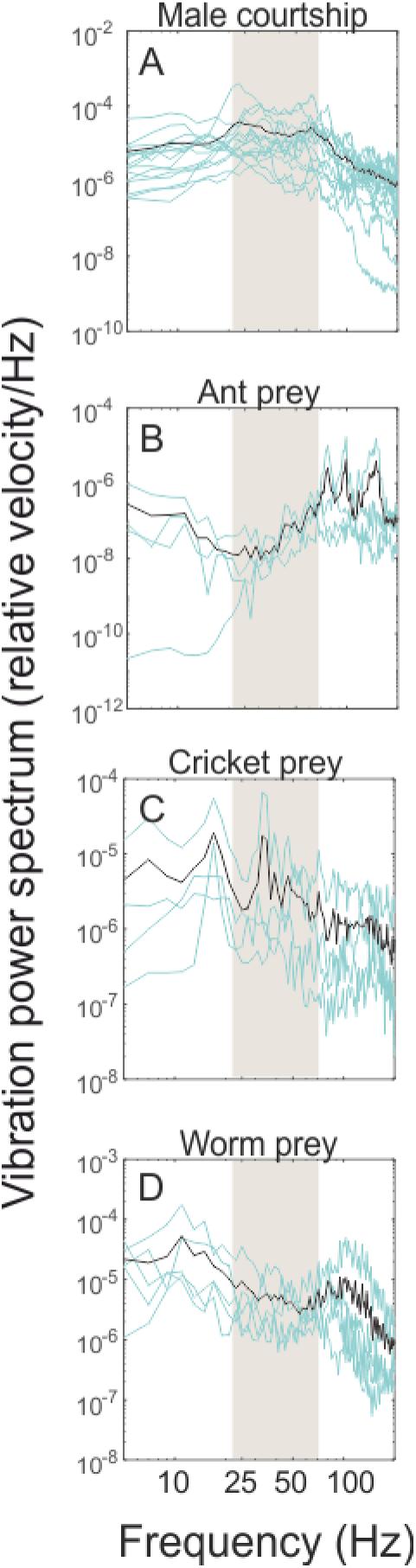
Male L. hesperus courtship and prey vibration power density spectra. (A)The courtship signals of males, who weigh 14 mg on average, mainly occupy the band between 23 and 61 Hz. The signals of certain potential prey items, which may differ in size such as (B) ants, (C) crickets (Acheta domesticus) and (D) mealworms (Tenebrio malitor) appear to reflect their mass, smaller prey items having higher frequencies and larger prey items having lower frequencies. In the species tested here, signals largely remain outside the frequency band occupied by males.

